# Antibiotic-mediated microbial community restructuring is dictated by variability in antibiotic susceptibility and population interactions

**DOI:** 10.1101/2024.08.18.608515

**Authors:** Kyeri Kim, Andrea Weiss, Helena R. Ma, Hye-In Son, Zhengqing Zhou, Lingchong You

**Affiliations:** Department of Biomedical Engineering, Duke University, USA; Center for Quantitative Biodesign, Duke University, USA; Center for Genomic and Computational Biology, Duke University, USA; Department of Molecular Genetics and Microbiology, Duke University School of Medicine, USA

## Abstract

It is widely known that faster-growing bacterial cells are more susceptible to antibiotics. Given this notion, it appears intuitive that antibiotic treatment would enrich slower-growing cells in a clonal population or slower-growing populations in a microbial community, which has been commonly observed. However, experimental observations also show the enrichment of faster-growing subpopulations under certain conditions. Does this apparent discrepancy suggest uniqueness about different growth environments or the role of additional confounding factors? If so, what could be the major determinant in antibiotic-mediated community restructuring? Combining modeling and quantitative measurements using a barcoded heterogeneous *E. coli* library, we show that the outcome of antibiotic-mediated community restructuring can be driven by two major factors. One is the variability among the clonal responses of different subpopulations to the antibiotic; the other is their interactions. Our results suggest the importance of quantitative measurements of antibiotic responses in individual clones in predicting community responses to antibiotics and addressing subpopulation interactions.

## Introduction

Environmental perturbations, such as temperature [1–3], nutrient [4], pH [5–7], and antibiotic treatment [8–12], can cause drastic perturbations to microbial community compositions and functionality. In the host gut microbiome, for example, antibiotic treatment leads the acute decrease of species richness as well as persistent alterations that were observed in taxonomy, metabolic output, and antibiotic resistance [13, 14]. More importantly, these alterations have a long-lasting impact in both adults and infants, raising risks to secondary infections and disease developments [15–19]. Therefore, many studies have examined the mechanisms underlying such restructuring dynamics in response to antibiotics [12, 20–22] and its functional implications [23]. A major focus of these studies is the presence of resistant species and their selection.

Even communities consisting of primarily antibiotic-suspectable species or strains exhibit highly complex responses upon antibiotic treatment. Pre-treatment microbiota structure can be one of the determinants of dynamics [14] but the varying phenotypic levels of antibiotic susceptibility among subpopulations can be the most significant factor that governs their survival dynamics [24]. Such heterogeneous resistance and cross-protection of sensitive populations change the community compositions [21, 22, 25–28]. Due to the complexity of the system, the mechanism is not clearly understood [12].

One of the most well-established mechanisms that drive population restructuring in antibiotic-sensitive populations is the positive correlation of growth and lysis rates [29–34]. In clonal populations, faster-growing bacteria are killed faster by antibiotics (Figure 1A). This notion has been widely demonstrated quantitatively in β-lactam treatment, where the killing rate is linearly correlated to the growth rate [30–34]. In single cells, this linear correlation can be explained by antibiotic-mediated filamentation: on average, faster-growing cells reach a critical length faster, leading to their faster lysis [33]. Given this explanation, one might in general expect the enrichment of a slow-growing population after antibiotic treatment, as shown with a significant fraction increase in the slow-growing subpopulation of *Salmonella* in mice [35].

**Figure 1:**
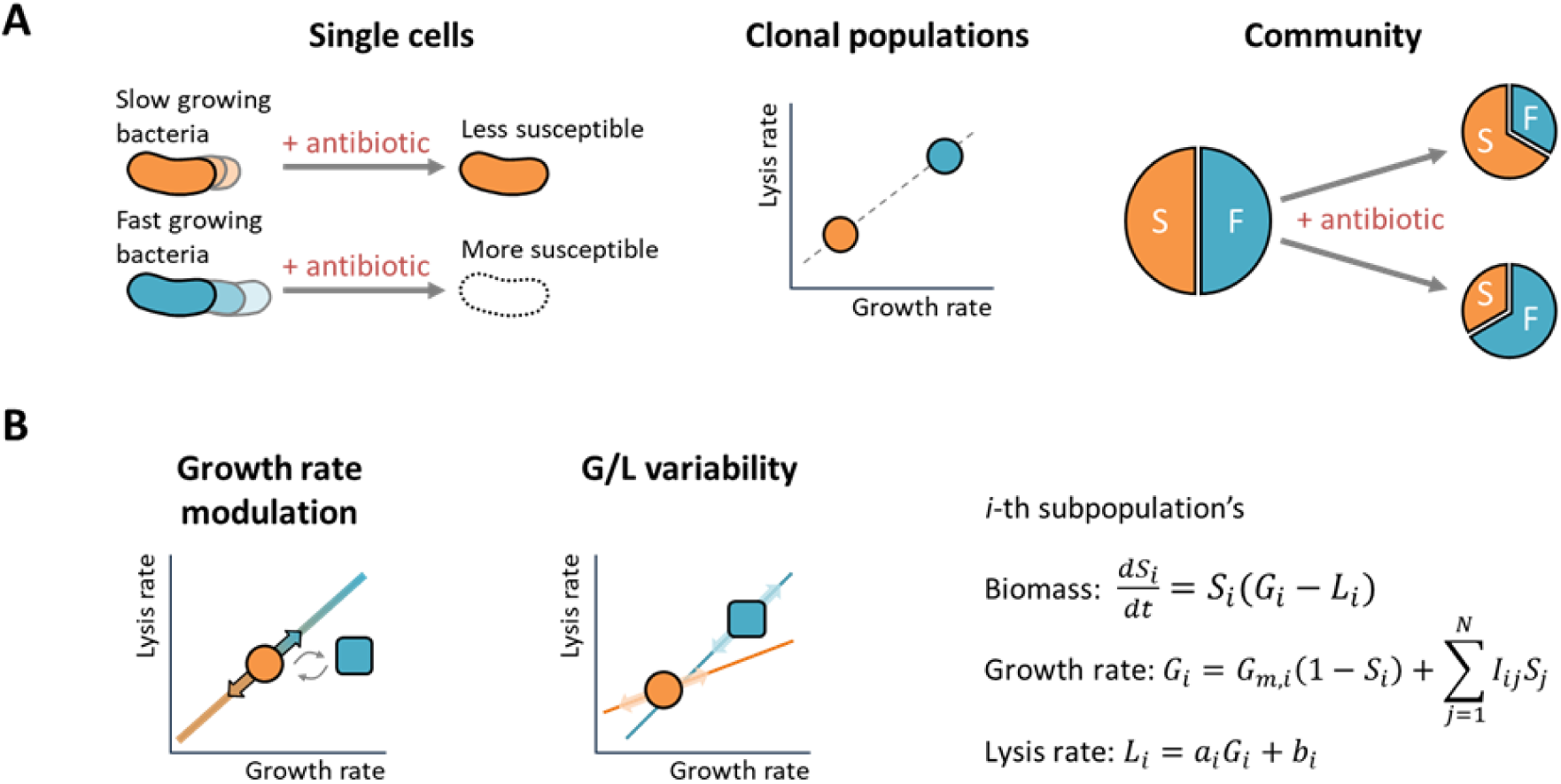
Enrichment dynamics in response to antibiotics. **A. Bacterial responses to antibiotic treatment from single cells to community dynamics.** In a single population, faster-growing cells are killed faster, leading to a positive linear correlation between the growth rate and the killing rate. Antibiotic treatment can lead to either an increase in the slow- or fast-growing fractions of the community, depending on growth and lysis rate correlations or subpopulation interactions, while concurrently reducing the overall population size. **B. Modeling the impact of interactions and variability of growth and lysis correlations on the community restructuring induced by antibiotics.** When a clonal population grows under community interactions or in different environments, changes of effective growth rate alter the lysis rate, along the linear correlation. Each subpopulation may follow a distinct linear correlation. Incorporating these two factors, subpopulation temporal dynamics can then be modeled. The abundance of i-th subpopulation (*S_i_*) increases and decreases with growth (*G_i_*) and lysis (*L_i_*) rates, where the *G_i_* accounts for logistic growth from the maximum growth rate (*G_m,i_*) and interactions (*I_ij_*) from *j*-th subpopulations. *L_i_* follows a unique, strain-specific linear correlation with *G_i_*, represented by the slope (*a_i_*) and intercept (*b_i_*).

In contrast to this intuitive expectation, studies have demonstrated the enrichment of fast-growing subpopulations during antibiotic treatment. In one example of a pair-wise community, where the ratio of the growth rates was larger than that of the lysis rates, Amor and Gore reported that a faster-growing *Corynebacterium ammoniagenes* population was enriched over a slower-growing *Lactobacillus plantarum* population *in vitro* [36].

However, in most cases, the linear correlation alone does not appear to explain the experimentally observed community dynamics. Schlomann *et al.* found that a faster-growing *Vibrio cholerae* population was enriched over a slower-growing *Enterobacter cloacae* population after sublethal antibiotic treatment in the zebrafish gut [37]. This enrichment was due to cell entanglement of the slow-growing subpopulation after antibiotic exposure and physical expulsion of aggregates from the host. In contrast, the mechanism is not always clear. For example, Certain *et al.* reported depletion of faster-growing cells *in vitro* (culture tubes) but their enrichment *in vivo* (mouse gut) [38]. This study suggested that potential intrinsic differences between antibiotic responses *in vitro* and *in vivo* led to diverging enrichment outcomes.

Here, we sought to determine the impact of two additional factors, variability in lysis kinetics and strain-to-strain interactions, on community composition dynamics. The variability in lysis kinetics indicates the variations in antibiotic susceptibility, with quantitative disparity in linear correlations among the strains and antibiotic conditions [31]. Interactions among strains can modulate subpopulation growth rates, directly affecting the effective lysis rates and leading to different composition outcomes [39].

We used a combination of mathematical modeling, high-resolution clonal measurements, and high-throughput NGS measurements of a barcoded library of single-gene knockout *E. coli* strains and found these two factors had predictable enrichment outcomes. Our results provide a coherent explanation for the diverse enrichment patterns reported in previous studies. Furthermore, our work underscores the value of comprehensively quantifying antibiotic responses across different bacterial species and strains, as well as interactions in a community.

## Results

### Modeling community response to antibiotic treatment

We modeled community restructuring by accounting for the growth and lysis of different members and their potential interactions, by using a generalized Lotka-Voterra (gLV) modeling framework (Figure 1B). The model consisted of a set of ordinary differential equations (ODEs) describing the abundance dynamics of subpopulations. The temporal dynamics of the *i-*th subpopulation, *S_i_*, are represented by the equation:

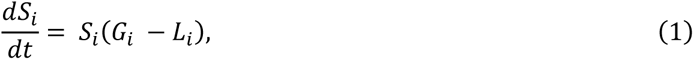

where *G_i_* and *L_i_* are the effective growth and lysis rates of the *i-*th subpopulation, which in general vary over time. We summarized the mathematical representations and definitions of the parameters used in the model in Table S1.

We assumed that *L_i_* is linearly correlated with *G_i_*:

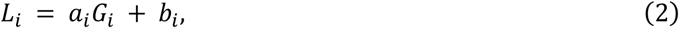

where *a_i_* and *b_i_* are the slope and intercept of the *i-*th subpopulation correlation, respectively. We termed this as G/L correlation.

For each subpopulation, *G_i_* is determined by the maximum growth rate (*G_m,i_,* which is a strain-specific intrinsic maximum rate that characterizes a growth of the subpopulation), the carrying capacity, and interactions with other subpopulations:

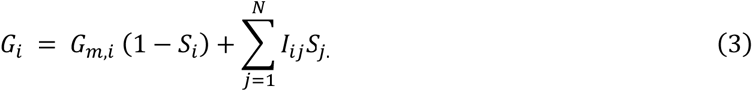

Here, *N* represents the total number of subpopulations. *I* is an interaction matrix. Each element, *I*_*ij*_, is a constant representing the uni-directional interaction from the *j*-th subpopulation to the *i*-th subpopulation with a sign (+/-) and its strength (magnitude). The diagonal elements of this interaction matrix are zero. The corresponding maximum lysis rate of *i-*th subpopulation *L*_*m*,*i*_ = *a*_*i*_*G*_*m*,*i*_ + *b*_*i*_. In sum, the biomass dynamics of the *i-*th subpopulation is described by (see the Methods section for detailed derivations):

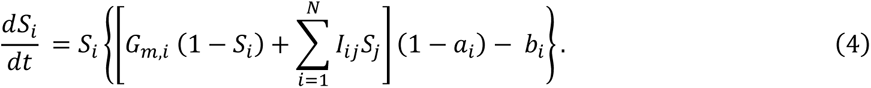

This model can be simulated with or without lysis, and with or without interactions among subpopulations. When no lysis occurs, *L*_*i*_ = *a*_*i*_ = *b*_*i*_ = 0. When no interaction occurs, *I* = *O* (zero matrix).

### Modeling predicts enrichment of slow-growing subpopulations in the absence of clonal variability in susceptibility and population interactions

Using the modeling framework, we simulated a community consisting of *N*(=30) subpopulations in two phases (both with the same duration, *T*): a growth-only phase (*L*_*i*_ = 0), where no antibiotic is added, followed by a phase involving both growth and lysis (*L*_*i*_ > 0), where an antibiotic is introduced (Figure 2A). The first panel displays the maximum growth (*G_m,i_*) and lysis (*L_m,i_*) rates of *i-*th subpopulation at a given condition as dot points. The solid line illustrates the linear G/L correlation of *i-*th subpopulation. It displays the corresponding *L_i_* when *G_i_* is modulated from *G_m,i_* by interactions. The maximum growth rates (*G_m_*) of the subpopulations ranges from 0.7 to 1.5. The marker color indicates *G_m_*, transitioning from orange to blue in increasing order of *G_m_*, while the precise gradient indicates the *G_m_* of each subpopulation. For the base case, two assumptions were applied. First, there is no growth rate modulation by interactions (i.e., *G*_*i*_ = *G*_*m*,*i*_ (1 − *S*_*i*_)). Second, all subpopulations follow the same linear G/L correlation: *L*_*i*_ = *aG*_*i*_ + *b* (where *a* = 1.5 and *b* = 0.1 in Figure 2A). The marker size indicates the net rate of each subpopulation without growth rate modulation (*G_m_ – L_m_*). The second panel indicates the net rates of the subpopulations, where the net rates of slower-growing subpopulations are larger. Therefore, in this base case, slower-growing subpopulations are less susceptible during the antibiotic treatment phase because the slope of the G/L correlation is larger than 1.

**Figure 2:**
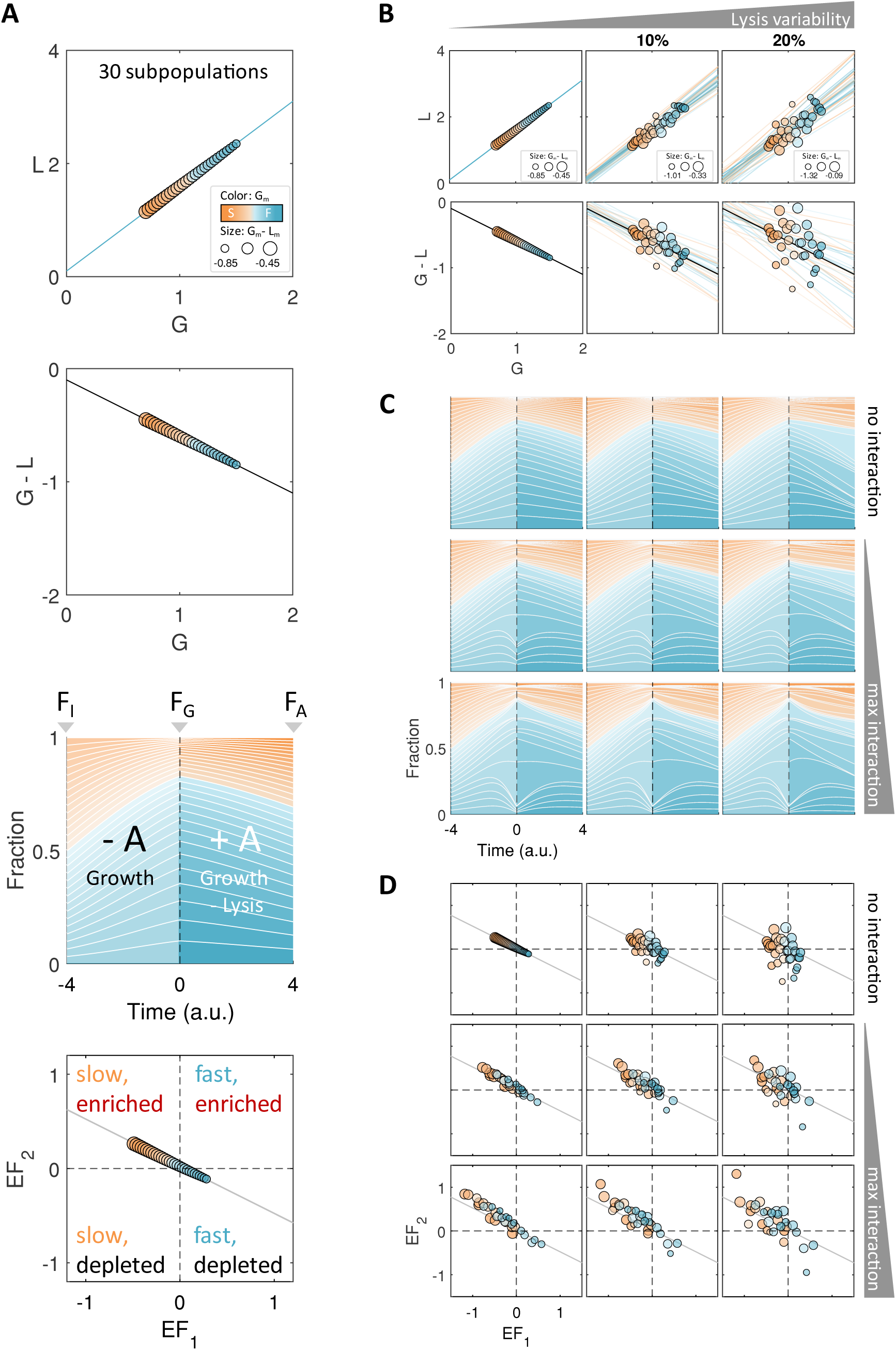
Subpopulation enrichment simulations incorporating growth-lysis linear correlation, variability, and random interactions. **A. Modeling illustrates the enrichment of slow-growing populations that have the same growth-lysis correlation and have no interactions beyond competition.** The first panel illustrates 30 distinct subpopulations assigned with varying maximum growth rates (*G_m_*) and corresponding maximum lysis rates (*L_m_*) which follows a unique linear correlation (*L = 1.5G + 0.1*). Markers are positioned by (*G_m_, L_m_*), colored by *G_m_*, and sized by the net rate (*G_m_ – L_m_*) of each subpopulation. The second panel shows the same lysis kinetics, but with y-axis in net rate (*G ‒ L*). The third panel shows the fractions of subpopulations over time in two phases: growth-only phase (-A) and a subsequent antibiotic-treated phase (+A), marked separation by a dashed line. *F_I_*, *F_G_,* and *F_A_* are the fractions at initial, after the growth, and after the antibiotic treatment phase, respectively. The fourth panel displays the enrichment factors (*EF*) of each subpopulation from two phases. Enrichment factors of the first growth phase (*EF_1_*) was calculated as 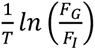 for each subpopulation and the second phase (*EF_2_*) was calculated as 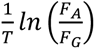, where *T* is the duration of phases. The solid line displays the baseline enrichment dynamics, where its slope is the same as lysis kinetics of the second panel, expected with no growth rate modulation (i.e., *G_i_* = *G_m,i_* at all times, without growth slow down, lysis rate variability, and interaction). In this base case, logistic growth slowdown was negligible, and therefore *EF*s of all strains were well aligned with the baseline enrichment dynamics. The faster-growing subpopulations (blue) resided towards the right side, and the greater net rate (*G_m,i_ - L_m,i_*) subpopulations (bigger markers) fell towards upper side. The four quadrants represent four qualitatively distinct enrichment outcomes (*EF_1_* > 0: relatively faster-growing, *EF_1_* < 0: relatively slower-growing, *EF_2_* > 0: enriched by antibiotic treatment, *EF_2_* < 0: depleted by antibiotic treatment). **B. Illustration of an increasing variability among the different populations in their G/L correlations. Starting** from a strict linear correlation of growth and lysis rates (G/L correlation) on the left, lysis rate variability was introduced by changing the slope and intercept of the G/L correlation to the right (10 and 20 % of the normally distributed random numbers were added, see Results and Methods section for detailed procedures). The G/L correlations of subpopulations are depicted as solid lines, showing the lysis dynamics with respect to the growth rate modulation (e.g., due to approaching its carrying capacity or interactions with other populations). The marker representation follows that of panel A. The second row shows the same lysis dynamics but represented as the net rate (*G-L*) over maximum growth rates (*G_m_*). **C. Fraction changes of subpopulations are shown over time when the simulations incorporate G/L variability and interactions.** To the base case (no variability and interactions) shown in top left and in panel A, G/L variability of corresponding degree in B was introduced (column-wise). Interaction was introduced upon the first row, shown in row 2 and row 3. The maximum interaction magnitude, max |*I_ij_*|, was set to 15 and 30, respectively. One random interaction matrix (*I*) was generated and shared by scaling up the elements. Therefore, the fraction changes in the first phases (-A, growth only) are same in each row. Different results of the fraction changes in the second phases are due to different lysis rates (column-wise), and interactions upon lysis (row-wise). **D. Antibiotic-mediated community restructuring is primarily driven by G/L correlations and further pushed by G/L variability and interactions among different subpopulations.** *EF*s of each phase were calculated from C. The color and size of the maker follows the same indicators in panel B. The linearly correlated antibiotic killing kinetics resulted in the linearly associated *EF*s of the community, close to the baseline enrichment dynamics. In increasing G/L variability, the *EF*s scattered from this tight linear association to all four quadrants, enable the diverse enrichment outcome. Increasing the magnitude of interaction terms (top to bottom rows) shuffled the order and broke the alignment of *G_m_* and net rate rankings to *EF_1_* and *EF_2_*, respectively, while the tight linear association of *EF*s remained. Combining two factors, *EF*s showed a shuffled, loose linear association spread from the baseline enrichment dynamics.

The third panel shows the simulated subpopulation fractions over time across the phases, starting from equal fractions (*1/N*) for all subpopulations. In the first phase, the fractions of subpopulations with larger *G_m_* (blue) mostly increased with relatively faster growth. In the second phase, the fractions of subpopulations with smaller *G_m_* (orange) increased due to their greater *G_m_ – L_m_*.

To illustrate the enrichment or depletion of each subpopulation, we quantified it by calculating the fold change per unit time in its fraction from the starting point to the end point, represented on a logarithmic scale. We term this metric the enrichment factor (*EF*). The *EF* of the *i-*th subpopulation during the first phase, 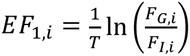, where *T* is the duration of the phase, *F* is the *i-*th subpopulation fraction at the end of the growth phase, and *F_I,i_* is its initial fraction. A subpopulation is enriched if its *EF* > 0 and depleted if *EF* < 0. If we assume each growth to be approximately exponential, *EF_1_* of the *i-*th subpopulation *EF*_1,*i*_ ≈ *G*_*m*,*i*_ + *c*_1_, where *c_1_* is a constant representation of the total population adjustment and is the same for all subpopulations (see Methods for further derivations). Likewise, in the second phase, 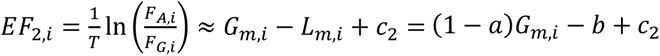, where *F_A,i_* is the fraction at the end of second phase and *c_2_* is a constant similar to *c_1_*. Therefore, *EF_1_* reports the extent of enrichment and is primarily dependent on *G_m_* during the growth phase; *EF_2_* is primarily dependent on net rate (*G_m_* ‒ *L_m_*) during the antibiotic phase.

The bottom panel of Figure 2A shows *EF_2_* plotted against *EF_1_*. Each dot represents one subpopulation, falling into one of four quadrants, each representing a unique enrichment pattern. In this base case, all points align with a straight line, termed as the baseline enrichment dynamics. It emerges from the linear G/L correlation, while its slope matches that of the second panel. The baseline enrichment dynamics were derived under the assumption that *G_m_* is constant in each subpopulation. Since the slope of the G/L correlation is greater than 1 (a>1), the negative linear *EF* association provides a quantitative depiction of the concept that fast-growing cells are killed faster. Consequently, most fast-growing subpopulations fall into quadrant 4 (bottom-right). Conversely, slower-growing subpopulations were depleted during the first phase but enriched in the second phase and fell into quadrant 2 (top-left). We note that faster-growing cells will be enriched during antibiotic treatment if *a* is less than 1, as reported by Amor and Gore [36].

### Modeling predicts diverse enrichment outcomes resulting from clonal variability in susceptibility and population interactions

Experimental evidence showed that both the slope and intercept of the linear G/L correlation are influenced by multiple factors, including antibiotic type, antibiotic dose, and bacterial species or strain [31–33]. In particular, under the same antibiotic treatment condition, the G/L correlations can vary among different subpopulations [31]. In addition, subpopulations often interact due to varied genetic backgrounds and growth conditions during antibiotic treatment [40–43]. We introduced these two factors into the model to examine their effects.

First, we introduced increasing degrees of variability to the base case G/L correlation between subpopulations (Figure 2B, top panels). The left-most panel represents the base case (Figure 2A). We generated two variability sets, by adding 10% or 20% of gaussian noise to the slope and intercept of the G/L correlations. The bottom panels show the net rates over the maximum growth rates. Figure 2C shows the simulated subpopulation fractions as a function of time, with corresponding variabilities (top row: no interactions). It shows that a high variability in G/L correlations led to substantial changes in the temporal dynamics of subpopulation fractions.

The increased variability in G/L correlations resulted in diverse enrichment outcomes (Figure 2D, top row). During the first phase, the enrichment was still determined by *G_m,i_*: subpopulations with larger *G_m,i_* had larger *EF_1,i_*. During the second phase, the *EF_2,i_* was determined by the net rate (*G_m,i_ - L_m,i_*). However, due to the variability in G/L correlations, a larger *G_m,i_* did not always correspond to a smaller net rate. The loss of this alignment caused the data points to scatter across all four quadrants. Some subpopulations with large *G_m,i_* were enriched during the second phase, while some subpopulations with small *G_m,i_* continued to be depleted. Therefore, the variability in the G/L correlation among subpopulations is one of the key factor contributing to the counter-intuitive enrichment of fast-growing populations.

Next, we introduced two increasing degrees of interactions into these variability configurations. A non-zero interaction matrix was generated by a previously published code [39], which assigns either positive and negative random numbers to the pairwise constant interaction rate (*I_ij_*) up to a specified interaction limit. The maximum magnitude of *I_ij_* was set to 10 times of the maximum *G_m,i_* for middle row and 20 times for the last row simulations (Figure 2C-D).

Compared to the effect of G/L variability alone, introducing interactions led to different changes in the enrichment pattern (compare Figure 2D top with the bottom two rows). The interactions can either promote or suppress the *G_i_*. In turn, *L_i_* is modulated indirectly, according to the strain-specific G/L correlation (Figure 1B, first panel). In the absence of variability in the G/L correlations (Figure 2D, left most panels in all rows), subpopulations with interactions still approximately followed a straight line, the baseline enrichment dynamics with small deviations. Moreover, the ranking of the subpopulations was not determined by *G_m,i_* but rather shuffled as a result of interactions. This could lead to the enrichment of the faster-growing subpopulations: subpopulations with larger *G_m,i_* grew more slowly due to interactions and became enriched during the antibiotic phase.

When both G/L variability and interactions affect community dynamics, we observe their combined effects on *EF.* The *EF* plot allows us to determine the primary factor influencing the diverse enrichment dynamics of each subpopulation. Deviations from the baseline enrichment dynamics are primarily due to the deviations from the average G/L correlation of the community. By incorporating clonal population growth and lysis rates onto the *EF* plot, we can reveal the influence of interaction and its consequences for enrichment dynamics.

### Experimental test of model predictions using a synthetic microbial community

To test model predictions, we assembled a synthetic community consisting of 73 strains from a barcoded Keio collection (Table S1), where the competition for nutrients (negative interaction) was the primary interaction within this community [39]. The Keio collection is a library of single-gene knockout strains derived from *E. coli* BW25113 ancestor strain [44]. In addition, each strain from our barcoded Keio collection contains a plasmid with a unique barcode sequence, enabling efficient community composition measurements using NGS [39, 45].

Figure 3 shows the workflow used to measure clonal population responses and community dynamics. Due to the single-gene knockouts, the fitness of each strain varies in the same growth environment. To confirm these fitness differences and collect growth and lysis dynamics for each strain, we inoculated 73 strains independently into glucose-supplemented M9CA medium with varying concentrations and types of antibiotics (Figure 3A, top row, and Figure S1; also see Methods for detailed protocols).

**Figure 3.**
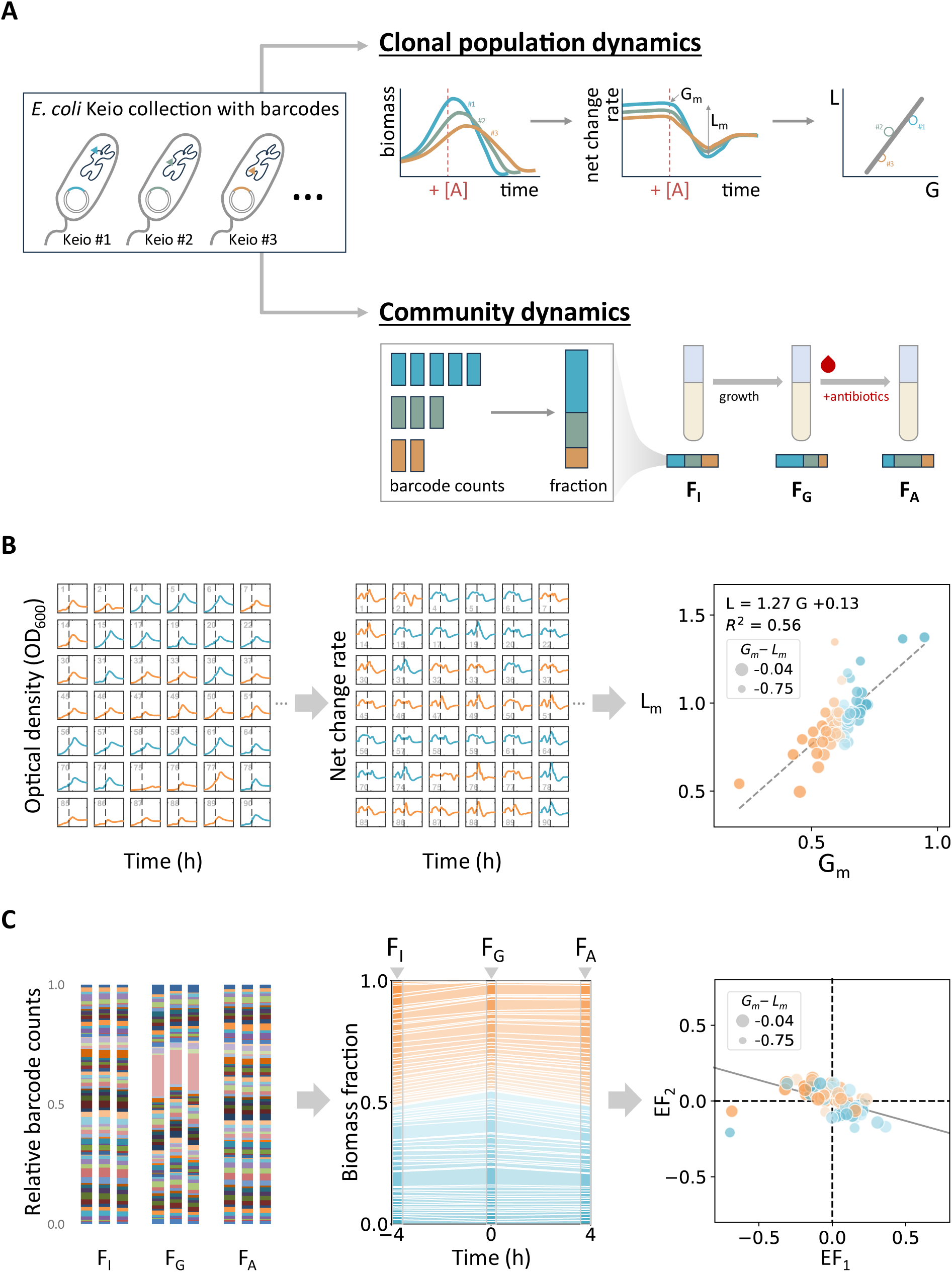
Quantifying antibiotic-mediated community restructuring in a synthetic microbial consortium. **A. The workflow of measuring clonal and community dynamics.** The community consisted of 73 *E.coli* strains from the single-gene knockout Keio library, which further barcoded by plasmids, each carrying a unique barcode sequence. Clonal populations were grown separately without antibiotics in the first phase, where the maximum growth rates were collected. Antibiotics were added subsequently. The maximum lysis rates were derived by the biggest difference in net change rates. G/L linear correlation was found by linear regression on 73 strain data points. An assembled community was grown without and with antibiotics through two phases and collected for composition measurements. Subpopulation fractions can be estimated by the relative barcode counts. Enrichment factors (*EF*s) were found from these fractions at 3 time point measurements: initial point (*F_I_*), after growth (*F_G_*), after antibiotic treatment (*F_A_*). **B. A linear G/L correlation was found from the measured clonal population dynamics.** In response to antibiotic treatment (5 µg/ml amoxicillin), clonal growth (*G_m_*) and lysis rates (*L_m_*) from these strains showed a linear correlation (dashed line displays L = 1.27G + 0.13) with substantial deviations from the common correlation in each strain (R^2^ = 0.56). Strains with smaller *G_m_* than the median of each community were colored orange, showing 36 slower-growing strains. The size of the marker corresponds to *G_m_ - L_m_* of each strain and the numerical indicator is shown in the same panel. **C. Antibiotic treatment led to complex community restructuring.** The relative barcode counts were collected from three technical repeats of the corresponding antibiotic condition at three time points: initial, after growth phase, and after antibiotic treatment phase. The median fraction was used for calibration to estimate biomass fraction (see Figure S2 for calibration details). *EF* from the first phase, 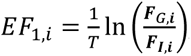, is shown in x-axis. *EF* from the second phase, 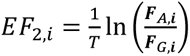 , is shown in y-axis. The marker representation follows that of panel B. The baseline enrichment dynamics (displayed in a solid line) was calculated from clonal population measurements in panel B, using Eq. 11 (see Methods). This community measurement shows that antibiotic treatment led to a general trend of slow-growing population enrichment (negative linear *EF* association), while data fell into all quadrants, scattered from the baseline enrichment dynamics. These reflect the linear G/L correlation and the variability in strains. Faster-growing strains (determined from panel B) were span through the entire *EF_1_* range. This reflects the presence of community interactions, modulation of growth rates.

In brief, we measured clonal population growth and lysis dynamics using time-course optical density at 600 nm (OD_600_) throughout the two phases: without and with antibiotics (Figure 3A-B). We then calculated the net change rate in biomass over time, profiling the maximum net change rate in the first phase as *G_m_* of each strain, which ranged from 0.2 to 1.0. For each antibiotic condition, we identified *L_m_* as the largest difference in net change rates from *G_m_.* In the condition with amoxicillin 5 µg/ml, *G_m_* and *L_m_* of the 73 barcoded Keio strains showed a linear correlation (L = 1.27G + 0.13, R^2^ = 0.56). The marker color and size follow the representation in Figure 2. Here, 36 subpopulations with *G_m_* smaller than the median of the community were labeled as slower-growing populations and depicted in orange color. The size of the marker corresponds to *G_m_* - *L_m_*, similar to the model simulation results and the numerical indicators displayed at the top left of the panel.

We assembled the community of all strains and measured compositions at 3 time points (Figure 3A bottom row): initially (*F_I_*), after the first phase (*F_G_*, growth-only), and after the second phase (*F_A_*, antibiotic-added). We measured compositions by utilizing next-generation sequencing of the barcode sequences. In brief, we sequenced three replicates of each condition (Figure 3C and S3A), calibrated them to biomass fraction (Figure S2), and calculated enrichment factors of each phase (Figure 3C).

We calibrated the relative barcode sequence counts using a mixture with known biomass fractions (Figure S2A). This calibration improved the estimation of biomass fractions: the calibrated fractions more closely followed the identity line (y=x) compared to the relative barcode counts without calibration. This calibration was tested with mixtures at varying biomass fractions from the same batch of overnight cultures (Figure S2B, rows 1 and 2). We also validated that this calibration was generalizable to the experimental replicates, using the community mixtures assembled from stationary-phase cultures and exponential-phase cultures from different batches of overnight cultures (Figure S2B, rows 3 and 4). In all 73 strains, the calibrated fractions closely aligned with the identity line (Figure S2C). Only two strains (ID 68 and 88) showed elevated calibrated fractions than the expected fractions in the small biomass fraction range (<0.001).

We visualized the enrichment dynamics in the same manner as presented in Figure 2, using the corresponding growth and lysis rates measured in panel B. The biomass fractions were ordered in increasing *G_m_* from top to bottom (Figure 3C, middle panel). The sum of fractions of the populations labeled as faster-growing from clonal population measurements (37 among 73 strains, marked in blue) were increased in the first phase but depleted during the second phase.

Consistent with model predictions (Figure 2D), the *EF*s of the given antibiotic condition showed an overall negative association, with significant deviations by individual populations from the baseline enrichment dynamics (Figure 3C, right panel) The baseline enrichment dynamics are displayed in solid lines, calculated using Eq. 11 in Methods, with parameters *a* and *b* from Figure 3B. Specifically, the enrichment plot shows that most of the populations labeled as faster-growing (marked in blue) were enriched in the first phase but depleted during the second phase, placing them in the bottom-right quadrant. In contrast, most of the populations labeled as slower growing (marked in orange) were depleted in the first phase but enriched in the second phase, illustrating canonical examples of slower-growing cells being enriched during antibiotic treatment. The deviation from the baseline enrichment dynamics (solid line) reflects the contribution of the variability in G/L correlations. In addition, the ranking of populations along the approximate direction of the baseline enrichment dynamics does not align with expectations based on the net rates (indicated by dot sizes), suggesting the contribution of population interactions.

### Diverse enrichment outcomes measured in the community dynamics across antibiotic conditions

We found that the overall linear G/L correlation remained across different β-lactam antibiotics treatments and concentrations (carbenicillin 20, 30, and 50 µg/ml, amoxicillin 3, 5, and 7 µg/ml, and cefotaxime 1, 2, and 4 µg/ml) (Figure 4A). These β-lactam antibiotics were picked based on the previous study demonstrating dose-dependent single-cell lysis dynamics [33] and linear G/L correlations [30, 31, 33]. In all cases, we observed deviations in data points from the overall G/L linear correlation to varying degrees. Corresponding antibiotic conditions showed linear *EF* associations (Figure 4B), consistent across four additional biological replicates without or with selection antibiotics (Figure S3C-D and S4). These linear *EF* associations reflect the general G/L linear correlations observed at the clonal population level.

**Figure 4:**
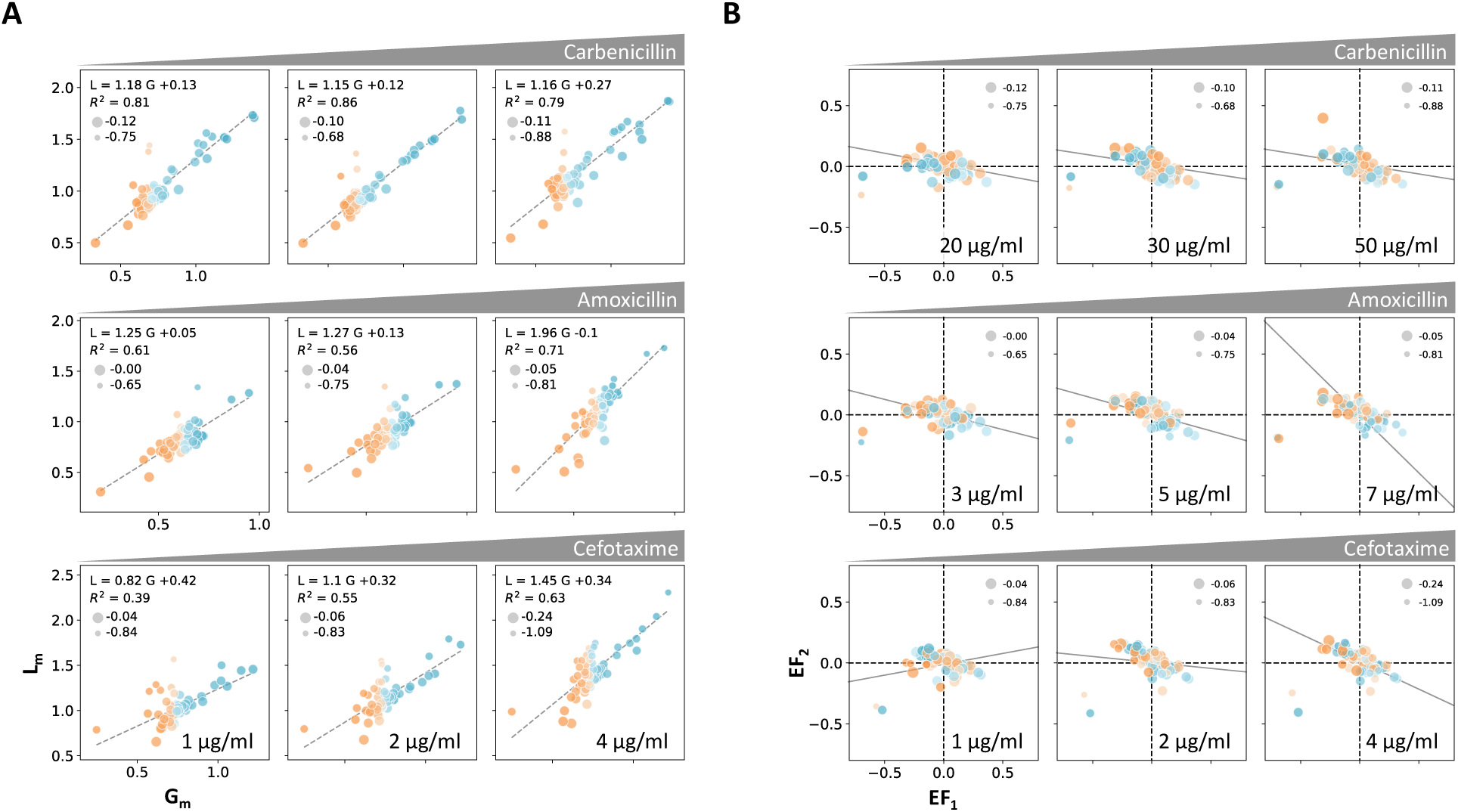
Linear G/L correlation in clonal population measurements and subpopulation enrichment at different antibiotic types and concentration studies. **A. The linear G/L correlation remained across the antibiotic types and concentrations.** The maximum growth rates and lysis rates (*G_m_* and *L_m_*) were determined from the clonal population growth and lysis curves, following the addition of carbenicillin (20, 30, and 50 µg/ml), amoxicillin (3, 5, and 7 µg/ml), and cefotaxime (1, 2, and 4 µg/ml). The linear regression of the 73-strain datapoints is depicted by dashed lines, with its equation and r-squared values displayed in the panel. The marker representation follows that of Figure 3B. **B. Enrichment of antibiotic treated community showed linear association.** The community coculture was treated with the corresponding antibiotic conditions in panels A, antibiotic was added after 4 hours of growth-only phase. Enrichment factors (*EF*s) were calculated by the endpoint-to-initial fraction ratios as shown in Figure 3C. Marker color and size representations follow those of panels A, while the numerical indicator size is also shown in the same panel. The baseline enrichment dynamics (displayed in a solid line) was calculated from the regression results shown in panel A (corresponding antibiotic conditions). In all conditions, datapoints are distributed from its baseline enrichment dynamics across all four quadrants, indicating the various enrichment outcomes, the linear G/L correlation, and presence of G/L variability. Strains spanned through the entire *EF_1_* without the order of *G_m_*, which reflects the presence of community interactions.

Importantly, the variability of G/L correlation was reflected in all antibiotic conditions by the spread in enrichment patterns from a tight linear association. The basal *EF* dynamic lines were derived from the regression of G/L correlation in corresponding antibiotic conditions shown in Figure 4A. These results support the diverse outcomes‒both qualitatively and quantitatively‒of enrichment dynamics.

We also observed negative *EF* associations in the non-treated community without additional dilution at the end of the first phase (Figure S3B, left panels). This correlation likely reflects growth approaching the carrying capacity. In particular, the measured OD_600_ of the non-treated community at the end of the second phase was higher than that of the overnight mixture (Figure S5). To further test this effect, we conducted simulations (Figure S6A). The results show that if faster-growing populations approach carrying capacity during the second phase, populations that are enriched during the first phase may have less room for further growth, leading to reduced enrichment or depletion. However, if populations experience approximately exponential growth in both phases, faster-growing populations will be more likely be enriched during the second phase as they were in the first phase. Further experiments recapitulated the predicted enrichment dynamics (Figure S6B): diluting the mixture from the first phase by more than 10-fold before starting the second phase resulted in an overall positive *EF* correlation between the two phases, as shown in the two right panels.

Considering the measured maximum growth rates (*G_m_*) and lysis rates (*L_m_*) of the clonal populations as *G_m,i_* and *L_m,i_*, we observed that the enrichment pattern resembled simulations incorporating interactions upon G/L variability. The alignments between *EF* and maximum rates (*EF_1_ : G_m_* and *EF_2_ : G_m_ - L_m_*) were partially mis-ordered. The experimentally determined Spearman correlation coefficients, which were less than 1, suggest the presence of interactions within the community. Perfect alignments would be indicated by a Spearman correlation coefficient of 1 (Table S3).

### Restructured population composition varied across different antibiotic types and conditions

Our community measurements showed that the final structure varied across different antibiotic types and concentrations, despite experiencing similar levels of population reduction in certain conditions (Figure 4B and Figure S5). Further analysis of the *EF_2_* data by strain (Figure 4B) revealed diverse enrichment results under different antibiotic conditions (Figure S7). While the direction of strain enrichment or depletion was consistent across antibiotic types for each strain, the extent of enrichment or depletion varied among antibiotic conditions. These results support that the positive linear G/L correlation impose the general trend of growth rate-dependent β-lactam killing kinetics. However, the exact G/L correlation, strain-to-strain variability on it, and interactions can influence its kinetics and sometimes shift and pivot the enrichment predictions.

Together, our results show that the antibiotic treatment imposes restructuring of the exposed community to varying final structures, which are majorly determined by growth and lysis dynamics of subpopulations in a given antibiotic condition and deviating further due to interactions between survivors.

## Discussion

Our findings underscore the importance of the two factors in explaining the diverse and sometimes paradoxical outcomes of antibiotic treatment observed in microbial communities [35–38]: the variability in growth and lysis kinetics of those clonal populations, and the community interactions among subpopulations.

Variations in antibiotic susceptibility depend on specific combinations of strains (or species), drug types, and drug concentrations. In the individual clonal killing curves of the single-gene knockout library treated with β-lactam antibiotics, our measurements showed that the linear G/L correlation is drug-specific, and the phenotypic variations in susceptibility from each correlation are evident among the strains. These results in a general trend of slower-growing population enrichment, yet with diverse enrichment patterns for individual strains. Under the β-lactam conditions that caused similar reductions in total population size, we observed the strain-specific enrichment varied across β-lactam types and concentrations. Since the survivor composition can influence the further dynamics and functionality [15–19], the ability to predict enrichment patterns can be valuable in designing antibiotic therapies and microbiome engineering strategies.

Considering that the G/L correlation can be mapped to the single-cell lysis kinetics [33], our results reveal that the connection from single-cell lysis kinetics to clonal population dynamics can be extended to community enrichment dynamics through G/L correlation. Thus, the enrichment factor association reflects experimentally measurable phenotypical traits at three levels of dynamics (single, clonal, and community), offering the quantitative mapping of bacterial response from single cells to community dynamics.

The effective growth rate of each member in a community often deviates from that of a clonal population [46]. This deviation is influenced by how subpopulations interact with each other and their environment. Our analysis leverages a library of *E. coli* strains with well-defined genetic backgrounds. Despite the similarity in their genetic backgrounds (aside from the single-gene knockouts), these strains exhibit diverse interactions, including both competition and cooperative interactions, depending on experimental conditions. In addition to demonstrating the dynamics predicted by our model, our analysis sets the stage for exploring the connections between genetics, phenotypic traits, interactions, and enrichment dynamics [47]. This capability can enhance our understanding of microbial responses to antibiotics and other environmental stimuli, ultimately contributing to more effective strategies for combatting antibiotic insusceptibility and optimizing treatment outcomes.

While our analysis focuses on β-lactams and a microbial community consisting of engineered *E. coli*, both our approach and conclusions can be broadly applicable. Our framework is adaptable to population responses of other classes of antibiotics, such as ribosome-targeting antibiotics, that exhibit growth rate-dependent susceptibility [48]. The general applicability is ensured by the coarse-grained nature of our framework. In particular, the lysis rate, ‘*L*’, in our framework can be interpreted as the rate of other forms of antibiotic-mediated death or growth suppression., and it can also be quantified by using the same pipeline. The framework is also applicable to communities consisting of diverse species and strains, even when some are resistant. The presence of both susceptible and resistant subpopulations essentially leads to highly divergent G/L correlations.

## Methods

### List of Bacterial strains, chemicals and reagents, and instruments

| Reagent/Resource | Reference or Source | Catalog Number |
| --- | --- | --- |
| **Experimental Models** |  |  |
| Barcoded *E. coli* Keio strains | Wang *et al*, 2022 |  |
| **Chemicals, enzymes, and other reagents** |  |  |
| M9 Medium Broth powder | VWR | J863 |
| Glucose | Acros organics | 388190010 |
| Thiamine hydrochloride | Sigma | T4625 |
| Magnesium sulfate | Sigma | M2643 |
| Calcium Chloride | OmniPur | 10035-04-8 |
| Bacto™ Casamino Acids, technical | Gibco | 223120 |
| Kanamycin sulfate | Sigma | 60615 |
| Chloramphenicol | Sigma | C0378 |
| Carbenicillin, Disodium Salt | Invitrogen | 10177-012 |
| Amoxicillin | Sigma | A8523 |
| Cefotaxime sodium salt | Sigma | C7039 |
| SealPlate Film, Sterile | Sigma | Z369667 |
| AeraSeal™ Film, Sterile | Sigma | A9224 |
| Phosphate buffered saline (PBS) | Sigma | P4417 |
| Q5 High-Fidelity 2X Master Mix | New England Biolabs | M0492 |
| Agarose | Bio-Rad | 1613101 |
| TAE Buffer (Tris-acetate-EDTA) (50X) | ThermoFisher | B49 |
| Zymoclean Gel DNA Recovery kit | Zymo Research | D4008 |
| Sodium hydroxide solution 2.0N | VWR | BDH7223-1 |
| Tris HCl (1 M), pH 7.0 | Quality Biological | 351-050-10 |
| MiniSeq Mid Output Kit (300 cycles) | Illumina | FC-420-1004 |
| **Software** |  |  |
| MATLAB R2021a |  |  |
| Python 3 |  |  |
| **Instruments** |  |  |
| Freedom EVO (automated liquid handling robot) | Tecan |  |
| Infinite 200 Pro (plate reader) | Tecan |  |
| Spark (plate reader) | Tecan |  |
| Thermocycler | Bio-Rad |  |
| MiniSeq | Illumina |  |

### Measurement of clonal population growth and lysis rates

Unless otherwise specified, barcoded Keio strains [39] were inoculated from glycerol stocks into fresh culture media in a deep well plate. The plate was sealed with AeraSeal film and incubated overnight (14-16 hours) in a 37°C plate shaker operating at 700 rpm. Each well contained 500 µl of M9CA media supplemented with 0.4% glucose, along with selection antibiotics (chloramphenicol 25 µg/ml and kanamycin 50 µg/ml).

Initial culture dilutions and antibiotic additions were set up by the Freedom EVO liquid handling robotic system. The operation included diluting overnight cultures into fresh media (M9CA with 0.4% glucose), mixing, and dispensing the diluted cultures into transparent, flat 384-well plates. These plates were manually sealed with transparent SealPlate film and incubated in a 37°C plate reader for continuous measurement of optical density at 600 nm (OD_600_, every 5 minutes), with a 5-second shaking before the measurements. Diluted β-lactam antibiotics were prepared from working stocks before addition to the media. Working stocks were prepared by dissolving antibiotic powders into molecular grade water and filtering (for carbenicillin and cefotaxime) or directly dissolving into dimethyl sulfoxide (DMSO, for amoxicillin). Antibiotics were then added to the 4-hour incubated cultures via robotic operation. Following antibiotic addition, the plates were resealed manually with a new transparent SealPlate film and returned to the 37°C plate reader.

To determine growth and lysis rates of the clonal barcoded Keio populations under each antibiotic condition, OD_600_ time-course data were analyzed using MATLAB. We adopted the methodology described in [31]. In brief, instantaneous net change rates over time were computed by determining the slope of linear regressions on log-transformed OD_600_, ln(OD_600_), within a 1-hour rolling window (comprising 12 data points) across the time-course measurements. The clonal population maximum growth rates (*G_m_*) were identified from the peak values of the first 4-hour instantaneous net change rates. The maximum lysis rates (*L_m_*) were derived from the difference between the maximum and minimum of the instantaneous net change rates (Figure S1).

### Calibration of relative barcode counts for estimating biomass fraction

To validate and enhance the accuracy of relative barcode counts as a surrogate of biomass fraction estimation, a comprehensive numerical calibration and its verification experiments were conducted (Figure S2). The biomass fraction was obtained from overnight cultures which were diluted to achieve an approximate OD_600_ range of 0.2 - 0.7 across all subpopulations, where OD_600_ is known to be linearly correlated with biomass. 73 subpopulations were then grouped into 7 tiers, representing varying biomass fractions within a mixture.

From the measured relative barcode counts obtained from an even volume mixture of all strains, a calibration vector was derived (Figure S2A). This vector comprises calibration factors for each strain, determined by the equation (biomass fraction in a mixed culture) = (calibration factor) x (measured relative count fraction). Here, the biomass fraction of the mixed culture was computed from individual OD_600_ measurements mentioned above. The measured relative count fraction was extracted from raw sequence count data, normalized by sum to 1.

To assess the efficacy of the calibration vector to biological repeats, mixtures of the 7 tier groups were prepared in different ratios, with a set of volume ratios selected to ensure relatively uniform distribution across a logarithmic scale (i.e., 10, 5, 1, 0.5, 0.1, 0.05, 0.02). Upon sequencing the permutation mix sets, the relative count fraction was calibrated by multiplying the calibration factors corresponding to each strain. The calibrated and non-calibrated measured fractions were compared by plotting them against the expected biomass fraction (see Figure S2B, first two rows respectively), with the identity line (y=x) shown as a solid line.

To ascertain the robustness of the calibration vector across biological replicates, the permutation mix sets from new overnight cultures and exponentially growing cultures were sequenced. The actively growing cultures were prepared from individually incubated subpopulations initiated at a 1:100 dilution to the fresh media. The calibrated fractions of the overnight and the actively growing replicate were compared with the expected biomass fractions (see Figure S2B, last two rows respectively). Further validation for each strain was shown (refer to Figure S2C). Notably, calibrated fractions from 71 strains exhibited excellent alignment with the identity line, while strains of Keio strain ID 68 and 88 displayed deviations, primarily in lower fraction ranges, from the identity line.

### Community barcode counts and enrichment calculation

A 73-strain community was generated by combining equal volumes of overnight cultures from the clonal populations. The mixture was 1:100 diluted to M9CA media containing 0.4% glucose, without selection markers. The diluted mixture was dispensed into a 96-deep well plate at 1 ml per well, sealed with AeraSeal film, and incubated in a 37°C plate shaker at 700 rpm. After 4 hours, β-lactam antibiotics were added to the respective conditions, and incubation continued at the same condition. At each time point (-4, 0, and 4 hours), 200 µl of the cultures were sampled for OD_600_ measurement, followed by centrifugation at 2000 g for 2-4 minutes (using the LSE Mini Microcentrifuge, Corning). The resulting pellets were resuspended in PBS, boiled at 95°C for 5 minutes using a thermocycler, and stored at 4°C until further barcode amplification, within 3 days.

For the preparation of the next-generation sequencing (NGS) library pool, established protocols as described by [39, 45] were followed. In brief, a short, 2-cycle PCR was performed to ligate dual indexes, sequencing primers, and adapters. Magnetic bead-mediated PCR product size selection was carried out. Sample pooling and amplification of the selected barcodes via a 30-cycle PCR followed. The final PCR products were identified by running a 2.5% agarose gel, with gel blocks sliced within the range of 190 – 290 bp. The amplified barcodes in the gel blocks were extracted using the Zymoclean Gel DNA Recovery kit. The final denatured library was prepared prior to sequencing in accordance with the manufacturer’s guidelines, with a 30% PhiX spike-in along with the library. Illumina MiniSeq and its associated software were utilized to perform pair-end sequencing, producing output into the fastq format. Barcode alignment and counting were performed by using a previously published Galaxy workflow of the established protocol. In brief, this workflow trims the barcode sequences between adapters and counts the number of aligned reads for each barcode of the strains, allowing for up to 2 bp-mismatches per read during barcode alignment.

A customized Python script via Jupyter Lab was employed for calculating and plotting the enrichment of subpopulations. Subpopulation fractions were determined by dividing the barcode counts by the total barcode counts in one community, followed by calibration through multiplication by a calibration vector. The enrichment of subpopulations between two points was computed as the log-scaled fraction ratio (fraction at the endpoint/ fraction at the initial point). Open-source code was used for strain-specific enrichment bar plots in circular form in Figure S7 [49].

### Model formulation and simulations

As discussed above, the model of community growth and lysis was formulated by a set of ordinary differential equations (ODEs), describing the biomass dynamics of each subpopulation. The derivation of Eq. 4 from Eqs. 1-3 is shown below:

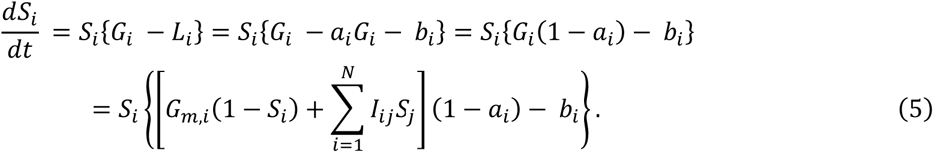

From the Eq. 1, when *G*_*i*_ remained *G*_*m*,*i*_ as a constant, *i-*th subpopulation fractions at initial time point, after growth, and after antibiotic treatment will be the followings, starting at the initial population size *S*_*i*,0_:

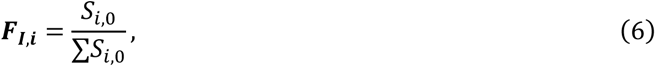

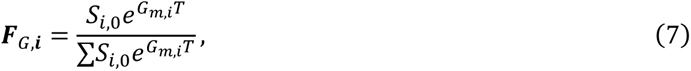

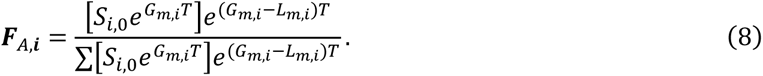

We defined enrichment factor (*EF*) as a ratio of fraction in logarithmic scale in each phase. Therefore, the first phase *EF* of the *i-*th subpopulation is the following equation:

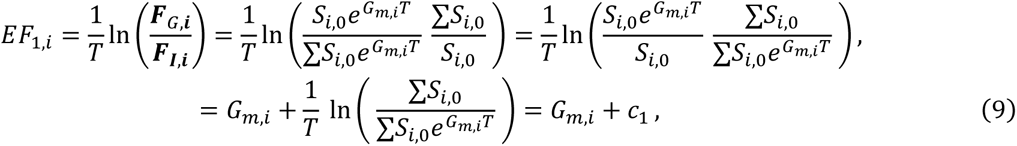

Similarly, the second phase *EF* of the *i-*th subpopulation is the following equation:

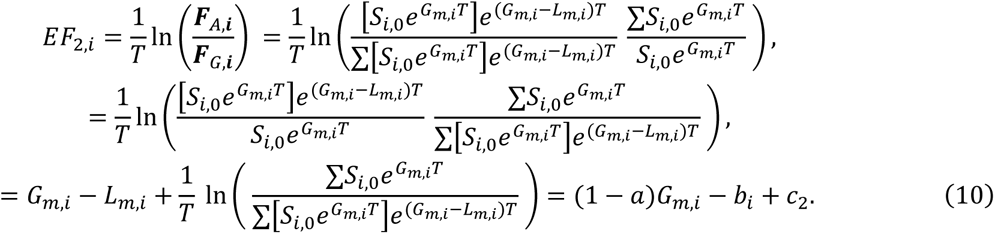

Therefore, when there is no G/L variability,

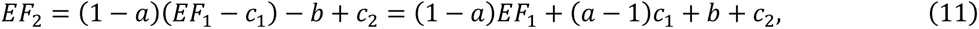

which reflects a linear association between enrichment factors.

For community formulation and simulations, the number of strains in a population, *N* was set to 30. Each subpopulation initiates with the same size of 0.01/*N* subpopulations at time *t* = - 4. Constant exponential growth rates (*G_m,i_*) were assigned to each subpopulation within the range of 0.7-1.5/hr. The lysis rates were assigned by a set of *a_i_* and *b_i_* to the *i-*th subpopulation. For no G/L variability, *a_i_* and *b_i_* were set to be constant (*a* and *b*). For x % of G/L variability, x % of a random number was added to the constants. That is, *a_i_* and *b_i_* were *a_i_* = *a* + x % random number, *b_i_* = *b* + x % random number. Simulations with variability share one random number set (30 random numbers were generated to have mean ∼0 and standard deviation ∼1). Fractions of each subpopulation at *t* = - 4, 0, and 4 (*F_I_, F_G_,* and *F_A_*) were retrieved for enrichment calculation. For non-treated population simulations with dilution factors, we resized the subpopulation by multiplying the dilution factor to all of the end point *S_i_* after the first phase simulation. All mathematical simulations were performed in MATLAB R2021a, with ode45 and ‘NonNegative’ option.

The interaction matrix was generated by the published code [39], which requires the number of subpopulations, ratio of the negative and positive interactions (50:50 was set in the community simulations), and maximum strength of negative and positive interactions as input of the code. The maximum positive interaction term was set to be half of the maximum negative interactions to 1) reflect the higher negative interactions observed in the same publication. The output of code is an N*-by-N* matrix where the pairwise interactions of *j*-th subpopulation to the *i*-th subpopulation are assigned to the elements *I_ij_*. Diagonal elements are 0. Note that simulations in Figure 2C-D share the same interaction matrix, the entire matrix was scaled to greater interaction strength.

## Acknowledgement

This work was partially supported by the National Institutes of Health (L.Y., R01AI125604, R01GM098642, and R01EB031869), US-Israel Binational Science Foundation (L.Y. 2021192), and the National Science Foundation (L.Y., MCB-1937259). The funders had no role in study design, data collection and analysis, decision to publish, or preparation of the manuscript.

