## Supplementary Information for "Antibiotic-mediated microbial community restructuring is dictated by variability in antibiotic susceptibility and population interactions"

### Table of contents

**Table S1 – S3** —— page 2-4

**Figures S1 – S7** —— page 5-13

| Parameter | Description |
|---|---|
| $\boldsymbol{G_i}$ | effective growth rate of i-th subpopulation |
| $\boldsymbol{G_{m,i}}$ | maximum growth rate of i-th subpopulation |
| $\boldsymbol{G_m}$ | maximum growth rates |
| $\boldsymbol{L_i}$ | effective lysis rate of i-th subpopulation |
| $\boldsymbol{L_{m,i}}$ | maximum lysis rate of i-th subpopulation |
| $\boldsymbol{L_m}$ | maximum growth rates |
| $\boldsymbol{G_i - L_i}$ | net rate of i-the subpopulation |
| $\boldsymbol{G_{m,i} - L_{m,i}}$ | net rate of i-the subpopulation without interactions |
| $\boldsymbol{a_i}$ | slope of the linear G/L correlation in i-th subpopulation |
| $\boldsymbol{b_i}$ | intercept of the linear G/L correlation in i-th subpopulation |
| $\boldsymbol{S_i}$ | size of the i-th subpopulation at time t |
| $\boldsymbol{I_{ij}}$ | Pairwise interaction rate from j to i-th subpopulation |
| $\boldsymbol{T}$ | Duration of each phase |
| $\boldsymbol{F_{I,i}, F_{G,i}, F_{A,i}}$ | Fraction of i-th subpopulation at the time point of initial, after first growth phase, after two (growth and antibiotic treated) phases. |

**Table S1. Model parameters for subpopulation growth and lysis model.**

| Keio ID | JW ID | Gene name | Keio ID | JW ID | Gene name |
|---|---|---|---|---|---|
| 1 | JW5419 | yfjP | 50 | JW5445 | ygcG |
| 2 | JW5438 | ygbF | 51 | JW5460 | ygeN |
| 4 | JW5481 | mltC | 52 | JW5532 | |
| 5 | JW5776 | sgcX | 53 | JW5803 | ybhR |
| 6 | JW5507 | yqiG | 54 | JW0689 | rhsC |
| 7 | JW0696 | ybfD | 55 | JW0323 | prpB |
| 8 | JW1282 | ycjD | 56 | JW0285 | yagX |
| 9 | JW2729 | ygcK | 57 | JW3386 | glpR |
| 10 | JW0026 | fkpB | 58 | JW3518 | tag |
| 11 | JW1464 | narU | 59 | JW5064 | |
| 12 | JW5198 | yciX | 61 | JW5428 | ygaY |
| 14 | JW5439 | | 64 | JW5577 | yigZ |
| 15 | JW5456 | ygeI | 65 | JW5142 | ycdR |
| 17 | JW5800 | nadR | 67 | JW1730 | ydjQ |
| 19 | JW1455 | ydcC | 68 | JW0345 | yaiL |
| 20 | JW1822 | yebS | 69 | JW1651 | ydhB |
| 22 | JW2875 | ubiH | 70 | JW3530 | glyS |
| 25 | JW5424 | ypjC | 74 | JW5451 | ppdB |
| 26 | JW5440 | ygcQ | 75 | JW5475 | rpiA |
| 27 | JW5457 | pbl | 76 | JW5646 | envC |
| 28 | JW5529 | yhbO | 77 | JW5432 | |
| 29 | JW5441 | ygcR | 78 | JW0486 | rhsD |
| 30 | JW3566 | rhsA | 80 | JW0820 | yliH |
| 31 | JW3451 | yhhI | 81 | JW3384 | rtcB |
| 32 | JW2528 | yphB | 82 | JW1572 | rspB |
| 33 | JW0746 | modA | 84 | JW2799 | mutH |
| 36 | JW2549 | recO | 85 | JW5437 | rpoS |
| 37 | JW5426 | yqaC | 86 | JW5454 | yqeH |
| 38 | JW5444 | ygcE | 87 | JW5476 | ygfI |
| 39 | JW5458 | ygeK | 88 | JW5707 | gspD |
| 40 | JW5531 | deaD | 89 | JW5442 | ygcU |
| 43 | JW0194 | metI | 90 | JW1451 | rhsE |
| 44 | JW0055 | yabP | 93 | JW3650 | yidF |
| 45 | JW3381 | malT | 94 | JW2505 | sseA |
| 46 | JW4056 | phnM | 95 | JW5183 | |
| 47 | JW4367 | thrL | 96 | JW2862 | xerD |
| 49 | JW5427 | ygaT | | | |

**Table S2. Keio strains included in the community coculture.**

| | | EF | Clonal rates | Correlation coefficients | | |
|---|---|---|---|---|---|---|
| | | | | Repeat 1 **(Figure 4B)** | Repeat 2 **(Figure S3C)** | Repeat 3 **(Figure S3D)** |
| **non-treated** ($t_0$) | | $EF_1$ | $G_m$ | 0.32 | 0.34 | 0.46 |
| **non-treated** (diluted at $t_0$) | | $EF_1$ | $G_m$ | 0.37 | 0.40 | 0.30 |
| **carbenicillin** | **20 μg/ml** | $EF_2$ | $G_m$ - $L_m$ | 0.16 | -0.06 | 0.33 |
| | **30 μg/ml** | $EF_2$ | $G_m$ - $L_m$ | -0.26 | -0.16 | 0.38 |
| | **50 μg/ml** | $EF_2$ | $G_m$ - $L_m$ | -0.04 | -0.10 | 0.56 |
| **amoxicillin** | **3 μg/ml** | $EF_2$ | $G_m$ - $L_m$ | -0.10 | -0.41 | -0.32 |
| | **5 μg/ml** | $EF_2$ | $G_m$ - $L_m$ | 0.07 | 0.08 | 0.41 |
| | **7 μg/ml** | $EF_2$ | $G_m$ - $L_m$ | 0.34 | 0.23 | 0.51 |
| **cefotaxime** | **1 μg/ml** | $EF_2$ | $G_m$ - $L_m$ | 0.30 | 0.11 | -0.10 |
| | **2 μg/ml** | $EF_2$ | $G_m$ - $L_m$ | 0.05 | 0.20 | -0.09 |
| | **4 μg/ml** | $EF_2$ | $G_m$ - $L_m$ | -0.13 | -0.12 | 0.12 |

**Table S3. Spearman correlation coefficients between G and enrichment by growth and between G-L and enrichment by antibiotics.**

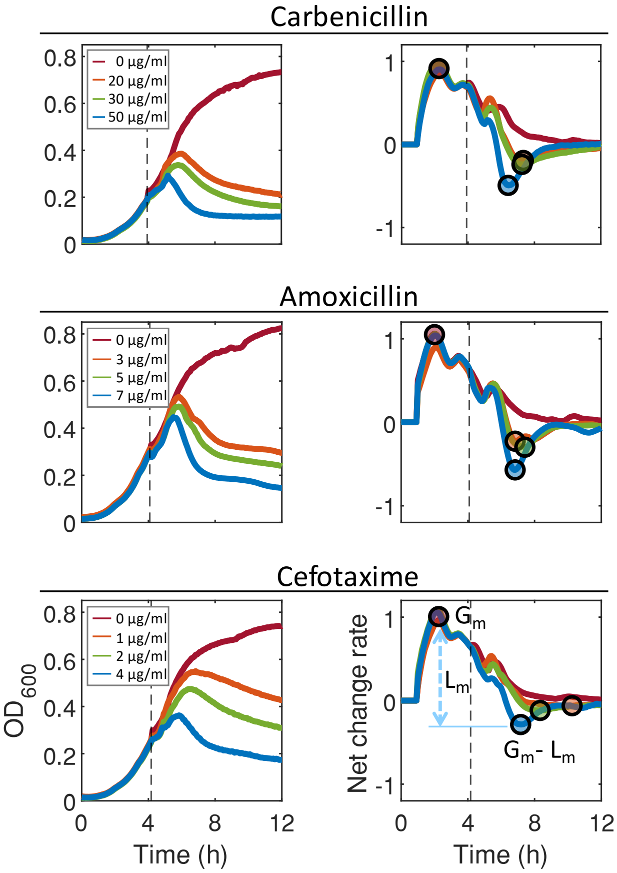


**Figure S1: Clonal population biomass curves and net change rates.**

OD time-course measurements of the Keio strain ID 95 in various antibiotic conditions are shown as an example for clonal population biomass measurements. Carbenicillin, amoxicillin, and cefotaxime were added to the culture with four different concentrations including non-treated control. Instantaneous net change rates were calculated from the OD measurements (see Method).

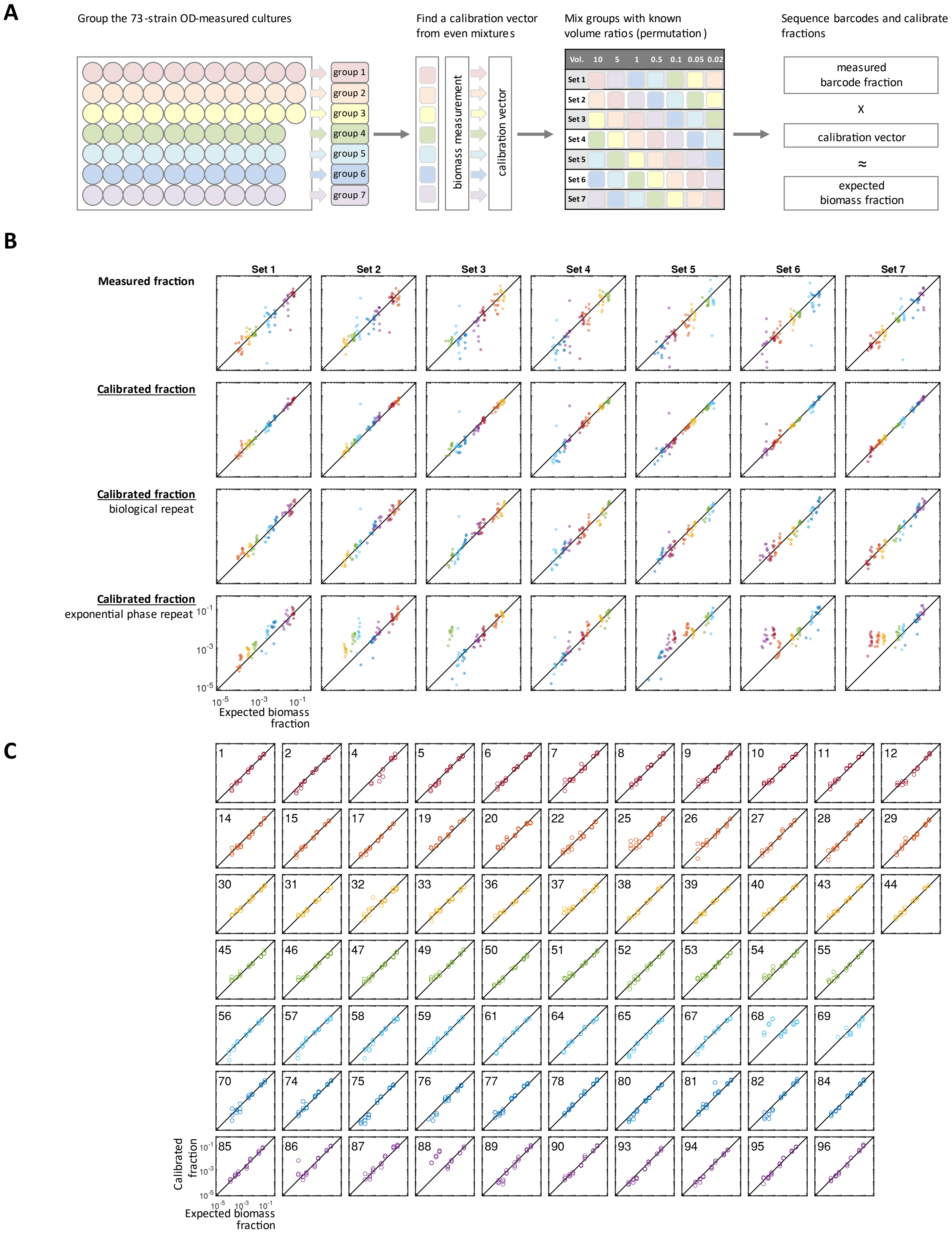

A
Group the 73-strain OD-measured cultures
group 1
group 2
group 3
group 4
group 5
group 6
group 7
Find a calibration vector from even mixtures
biomass measurement
calibration vector
Mix groups with known volume ratios (permutation)
Vol. 10 5 1 0.5 0.1 0.05 0.02
Set 1
Set 2
Set 3
Set 4
Set 5
Set 6
Set 7
Sequence barcodes and calibrate fractions
measured barcode fraction
x
calibration vector
≈
expected biomass fraction
B
Set 1 Set 2 Set 3 Set 4 Set 5 Set 6 Set 7
Measured fraction
Calibrated fraction
Calibrated fraction
biological repeat
Calibrated fraction
exponential phase repeat
Expected biomass fraction
C
1 2 4 5 6 7 8 9 10 11 12
14 15 17 19 20 22 25 26 27 28 29
30 31 32 33 36 37 38 39 40 43 44
45 46 47 49 50 51 52 53 54 55
56 57 58 59 61 64 65 67 68 69
70 74 75 76 77 78 80 81 82 84
85 86 87 88 89 90 93 94 95 96
Calibrated fraction
Expected biomass fraction

**Figure S2: Numerical calibration of the measured barcode count fraction improved to indicate the expected biomass fraction.**

A. **Illustration of the workflow of finding numerical calibration vector and its validation.** $OD_{600}$ of the diluted overnight culture were measured. From even volume ratio, calibration vector was derived by dividing expected biomass fraction by relative barcode count fraction. Calibration vector's applicability was tested with known logarithmic volume ratio distribution (10, 5, 1, 0.5, 0.1, 0.05, 0.02), while 73 subpopulations were grouped into 7 tiers of each permuting volume tier (See Method for detailed protocol).

B. **Robustness of calibration in the estimation of biomass fraction across the biological repeats.** The non-calibrated (first row), calibrated (second row), calibrated biological repeat (third row), calibrated repeat with actively growing strains (forth row) fractions were plotted over the expected biomass fraction. The identity line (y=x) is shown as a solid line (See Method for detailed protocol).

C. **Strain-specific calibration results.** Calibrated fraction for each strain was shown. Among 73 strains, 71 strains exhibited great alignment with the identity line, while strains of Keio strain ID 68 and 88 showed deviated calibrated fraction in lower biomass fraction range.

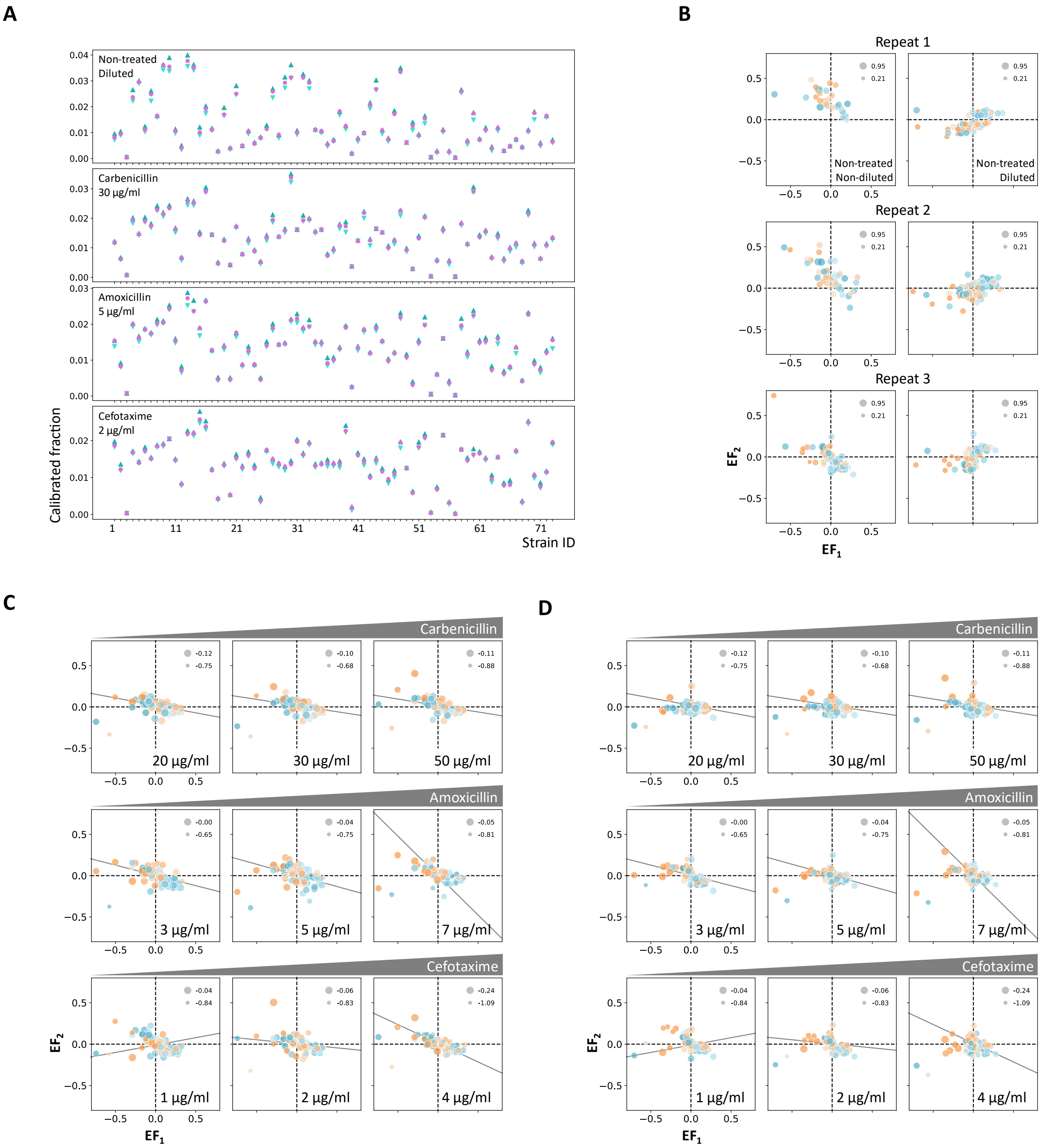


**Figure S3: Biological repeats and technical replications for the enrichment measurements**

**A. The median of the 3 technical replicates was selected for enrichment analysis**. Each biological repeat comprised 3 technical replicates (independent growth, antibiotic addition, and sampling) for a given experimental condition. The maximum (upward triangle, dark blue), median (circle, pink), and minimum (downward triangle, light blue) of the calibrated fractions of those replicates were shown in each strain. 4 representative experimental conditions (displayed in each plot) were selected for the presentation.

B. **Enrichment of the non-treated populations in biological repeats.** Biological repeats were placed row-wise. In each row, the left panel shows non-diluted community which corresponds to the "No dilution" condition in Figure S6 and 0 µg/ml control (in red) in Figure S5. The right panel shows the diluted (1:25) community which corresponds to the "1/25" conditions in Figure S6 and 0 µg/ml-diluted control (in pink) in Figure S5.

**C-D. Biological repeats of enrichment measurements.** Two biological repeats of the enrichment measurement were presented, apart from Figure 4B. These repeats were conducted with new batches of overnight cultures in combining strains into a community.

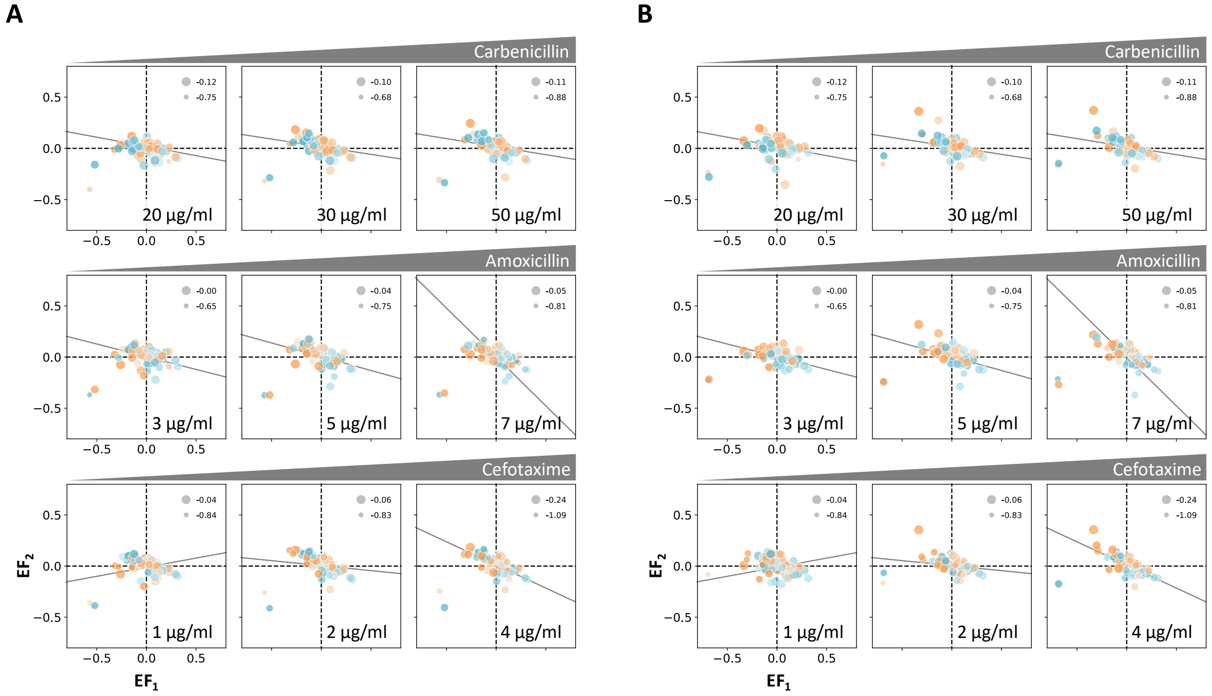


**Figure S4: Biological repeats with selection antibiotics.**

Two more biological repeats of Figure 4B with selection antibiotics were performed. Results depicted in panel **A** used the media containing chloramphenicol (25 µg/m) which selects for barcode-containing cells. Results shown in panel **B** used the media containing both chloramphenicol (25 µg/ml) and kanamycin (50 µg/ml) which selects Keio strains. These repeats were conducted with the same initial community as Figure 4B.

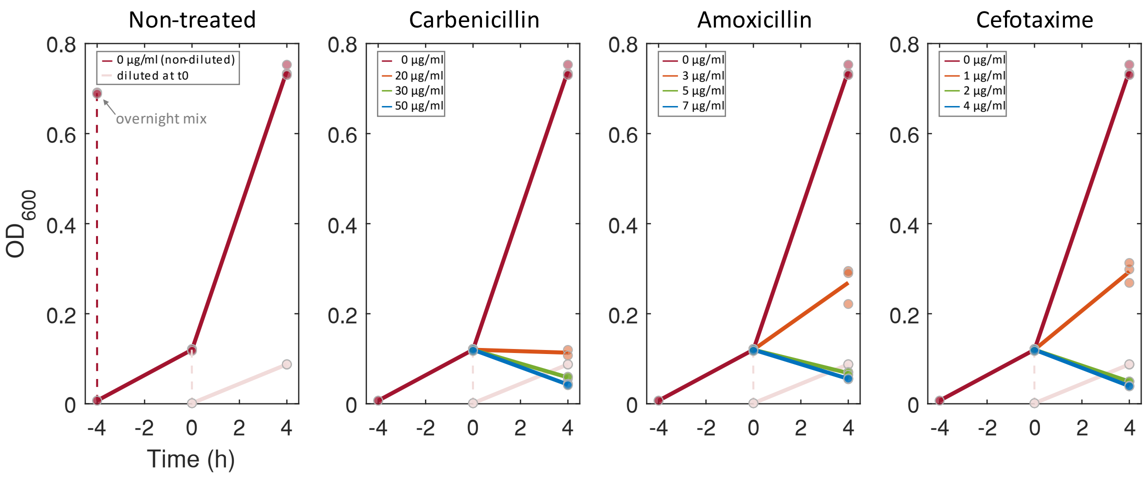


**Figure S5: Total population sizes after the growth and antibiotic treatment phases.**

The average $OD_{600}$ measurements (subtracted by the blank media) of non-treated, non-treated and diluted, and antibiotic-treated communities are shown in solid line. Technical repeat measurements were shown in dots. Note that 0 µg/ml control (red) was diluted (1:25) at time 0 to the fresh media to maintain the small population size and exponential growth (pink).

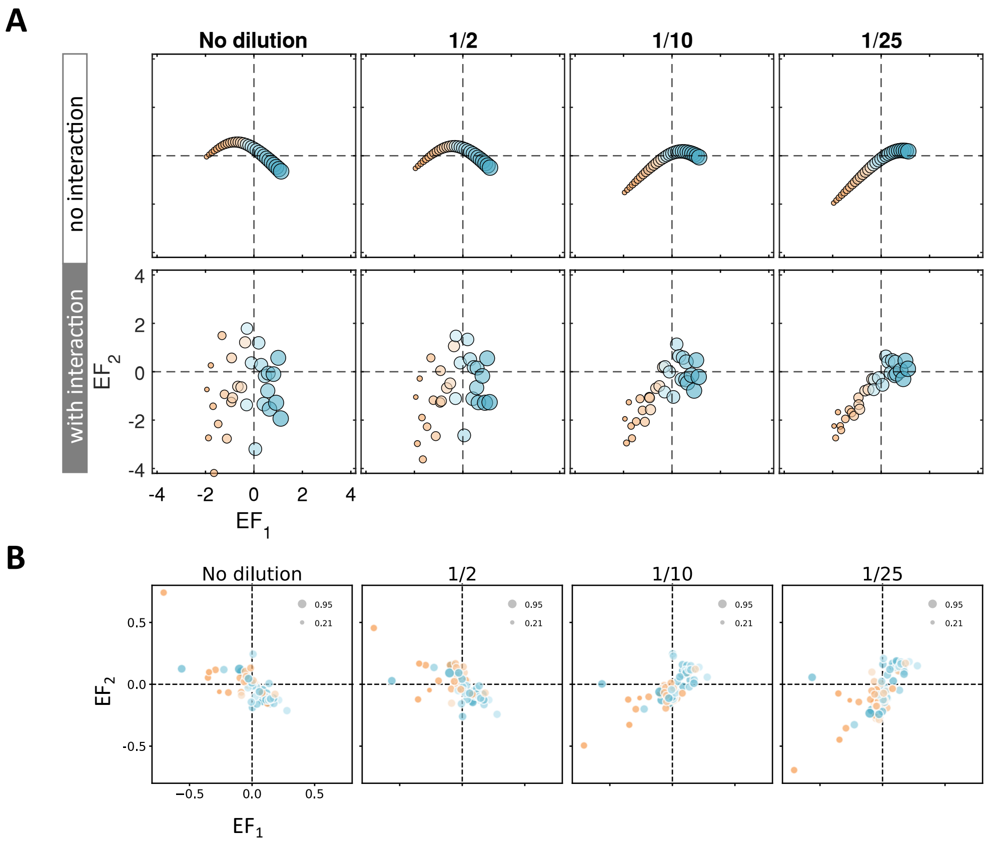


**Figure S6: Enrichment of the non-treated community with decreasing dilution ratio.**

A. **Simulations of non-antibiotic treated community enrichment results.** No interaction was considered in the first row. Maximum interaction factor, 1, was introduced to this model. Subpopulations with larger $G_m$ than the median of the community were marked in blue. The size of the markers corresponds to $G_m$ due to no lysis in the second phase. The numerical indicator is shown in each panel.

B. **Experimentally measured enrichment in corresponding dilution conditions.** Enrichment of 4 hours of the first growth phase is shown in $EF_1$. The culture was subjected to further dilutions to fresh media with 1, 1/2, 1/10, 1/25 dilution factors. Enrichment of another 4 hours of growth phase is shown in $EF_2$. Positive enrichment correlation with greater dilution implies that maintaining a small total population size enables exponential-like growth, by reducing the logistic growth term in the model.

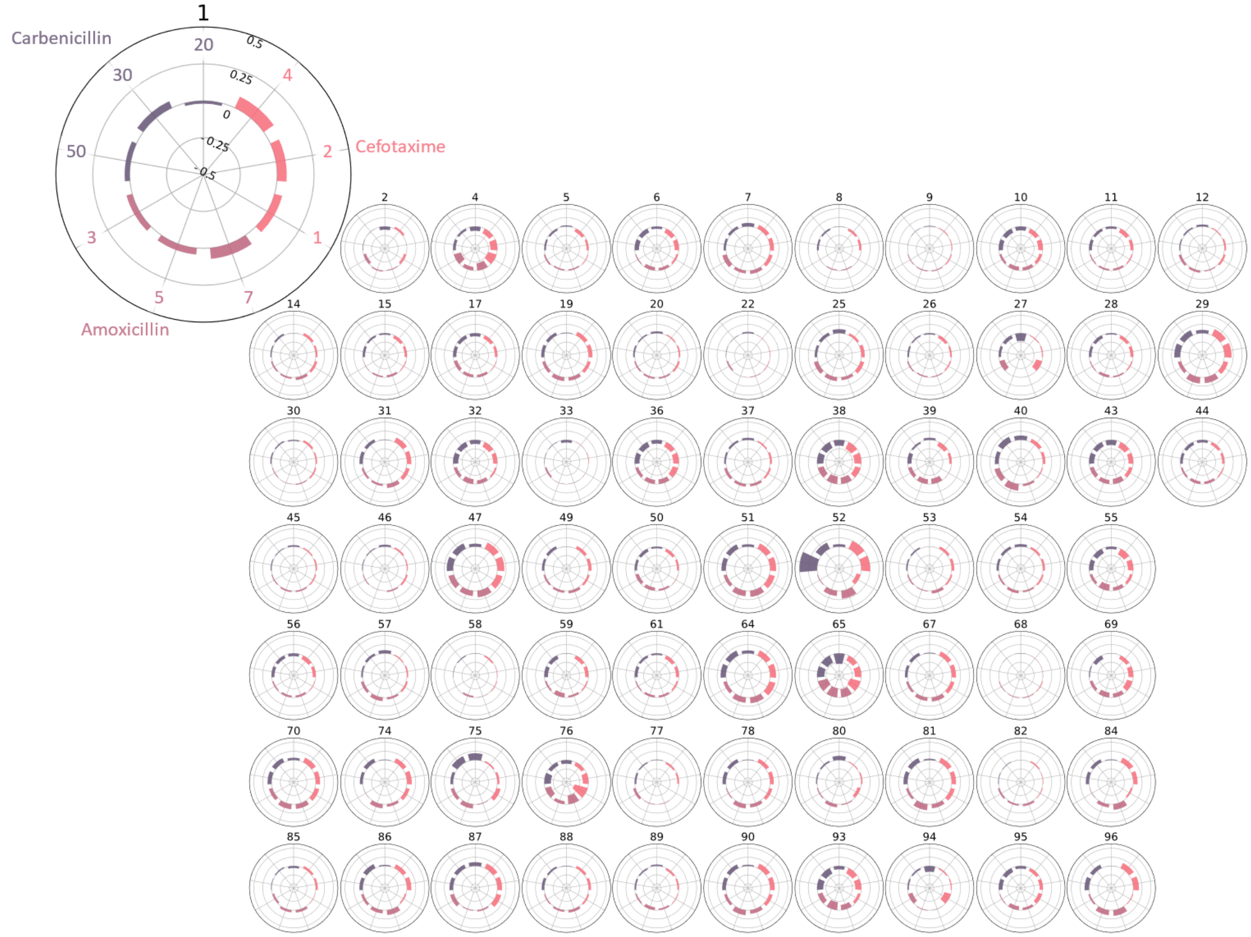


**Figure S7: Strain-specific antibiotic-induced enrichment comparison in the tested conditions.**

The $EF_2$ (Enrichment Factor of the antibiotic treatment phase) shown in Figure 4B is depicted by strain as circular bars. The bar chart starts at 0, referring to no fraction changes, and ranges from -0.5 to 0.5 from the center to the exterior of the circle. Antibiotic types and concentrations are displayed in corresponding colors: carbenicillin (20, 30, and 50 µg/ml) in purple, amoxicillin (3, 5, and 7 µg/ml) in violet, and cefotaxime (1, 2, and 4 µg/ml) in pink. Corresponding strain ID number is shown as the title of each panel.
